# Sulfoquinovosyl diacylglycerol is required for dimerization of the *Rhodobacter sphaeroides* RC-LH1 core complex

**DOI:** 10.1101/2024.03.24.586425

**Authors:** Elizabeth C. Martin, Adam G.M. Bowie, Taylor Wellfare Reid, C. Neil Hunter, Andrew Hitchcock, David J.K. Swainsbury

**Affiliations:** Plants, Photosynthesis and Soil, School of Bioscience, University of Sheffield, Sheffield, UK; School of Biological Sciences, University of East Anglia, Norwich, UK; Centre for Bacterial Cell Biology, Newcastle University, Newcastle, UK

## Abstract

The reaction centre-light harvesting 1 (RC-LH1) core complex is indispensable for anoxygenic photosynthesis. In the purple bacterium *Rhodobacter* (*Rba.*) *sphaeroides* RC-LH1 is produced both as a monomer in which 14 LH1 subunits form a crescent-shaped antenna around one RC, and as a dimer, where 28 LH1 subunits form an S-shaped antenna surrounding two RCs. The PufX polypeptide augments the five RC and LH subunits, and in addition to providing an interface for dimerization, PufX also prevents LH1 ring closure, introducing a channel for quinone exchange that is essential for photoheterotrophic growth. Structures of *Rba. sphaeroides* RC-LH1 complexes revealed several new components; protein-Y, which helps to form a quinone channel; protein-Z, of unknown function but which is unique to dimers; and a tightly bound sulfoquinovosyl diacylglycerol (SQDG) lipid that interacts with two PufX arginines. This lipid lies at the dimer interface alongside weak density for a second molecule, previously proposed to be an ornithine lipid. In this work we have generated strains of *Rba. sphaeroides* lacking protein-Y, protein-Z, SQDG or ornithine lipids to assess the roles of these previously unknown components in the assembly and activity of RC-LH1. We show that whilst the removal of either protein-Y, protein-Z or ornithine lipids has only subtle effects, SQDG is essential for the formation of RC-LH1 dimers but its absence has no functional effect on the monomeric complex.

## Introduction

The purple phototrophic bacterium *Rhodobacter* (*Rba.*) *sphaeroides* contains hundreds of specialised chromatophore vesicles, around 50 nm in diameter, the numbers of which respond to the incident light intensity [1–3]. The major membrane complexes of the chromatophore comprise the reaction centre-light harvesting 1 (RC-LH1) core complex, the peripheral light harvesting 2 (LH2) antenna complex, the cytochrome (cyt) *bc*_1_ complex, and ATP synthase [4]. Of these, cryogenic electron microscopy (cryo-EM) structures of cyt *bc*_1_ [5], LH2 [6] and both the monomeric and dimeric forms of RC-LH1 [7–10] have recently been determined. Within the chromatophore, LH2 complexes each bind a circular array of bacteriochlorophyll (BChl) and carotenoid pigments that absorb light and transfer the energy via the LH1 complex to the RC, where the energy is transiently stored as a charge separation [11–13]. Light-driven reduction of quinone to quinol at the RC is followed by passage of quinol across the LH1 ring [13] and diffusion to a cyt *bc*_1_ complex, where the quinol is oxidised through operation of the Q-cycle [14–16]. The cyt *bc*_1_ complexes reside in locally lipid-rich domains [4,17] located within a few nm of the RC-LH1 complexes [18]. The catalytic mechanism of the cyt *bc*_1_ complex reduces cytochrome *c*_2_ [11,19], which returns to the photo-oxidised RC, completing the cyclic ET pathway [11,20]; the Q-cycle mechanism also releases protons into the chromatophore lumen, generating a proton-motive force that is utilised by ATP synthase to drive the production of ATP to power cellular metabolism [20,21].

In *Rba. sphaeroides*, the RC is comprised of 3 subunits (L, H and M) and is surrounded in a fixed stoichiometry by a crescent-shaped LH1 ring containing 14 pairs of α and β transmembrane polypeptides, each of which binds two BChls and two carotenoids [7–10,22,23]. Closure of the LH1 ring is prevented by a single copy of the PufX subunit, which precludes the insertion of further LH1 subunits to maintain the open complex [7–10,24–28]. One copy of the recently discovered protein-Y subunit (using the naming convention suggested by Swainsbury *et al* [29]) sits between the RC and the LH1 ring, creating the RC_3_-LH1_14_-XY complex and maintaining a separation that allows quinones and quinols to diffuse freely to and from the RC Q_B_ site.

In purple bacteria most RC-LH1 complexes are monomeric, some with a closed LH1 ring as in *Rhodospirillum rubrum* [30] and *Thermochromatium* (*Tch*.) *tepidum* [31] and others with an incomplete ring - held open by extra scaffolding subunits such as PufX (*Rba. veldkampii*) [32], protein W (*Rhodopseudomonas palustris*) [33–35], or by the insertion of a transmembrane helix associated with the cytochrome subunit (*Roseiflexus castenholzii*) [36,37]. The many structural variations of RC-LH1 complexes are reviewed in [13]. In this context, RC-LH1 complexes in *Rba. sphaeroides* are unusual because they form dimers in which 28 LH1 subunits form an S-shaped assembly around two RCs, which creates a seamless path for energy transfer between the two halves of the complex. This arrangement provides an elegant energy-conservation mechanism that allows an LH1 excited state access to a second RC if the first is already undergoing a charge separation [38,39]. In addition, the dimeric complex is bent with the two monomers held at an angle of 152° [7–10], which imposes curvature on the membranes and is partly responsible for the spherical shape of the chromatophore vesicles [1,8–10,23,40–43]. This property is unique to a few close relatives of *Rba. sphaeroides,* which were recently reclassified into the *Cereibacter* genus [44]. To ensure consistency with the previous literature we will refer to these species as belonging to the *Cereibacter* subgroup and continue to use the historical species names for *Rba. sphaeroides* and its relatives throughout this manuscript. Mutants of *Rba. sphaeroides* that lack PufX are not only unable to grow photoheterotrophically but no longer form dimers [17,32–35,45,46].

Recent cryo-EM structures of the monomeric and dimeric forms of *Rba. sphaeroides* RC-LH1 have revealed further components in the complex beyond PufX. Protein-Y forms a hydrophobic hairpin structure that lies between the inside surface of LH1α 13 and 14 and the RC. Genetic removal of protein-Y (also called protein U) results in the formation of an incomplete LH1 ring with as few as 11 α and 10 β subunits [9,10]. This suggests that in addition to promoting the access of quinones to the RC Q_B_ site, protein-Y also provides a binding site for the LH1 subunits at the edge of the LH1 array, ensuring a gap that is correctly positioned to facilitate rapid diffusion of quinone and quinol between the RC and the external quinone pool. Four copies of a second novel transmembrane polypeptide, protein-Z, were also identified in the dimeric structure, but are absent in the monomer, suggesting an as yet undetermined role unique to the dimer [8]. The dimeric structure also revealed the presence of two lipids, the first of which was confidently assigned as sulfoquinovosyl diacylglycerol (SQDG) based on clear density of its distinctly shaped sulfonic acid head group. The other lipid was less well defined but was proposed to be an ornithine lipid [8]. The SQDG lipid was found to bridge the two halves of the dimer, interacting with both the RC L subunit and the critical Arg49 and Arg53 residues of PufX in one monomer, and an LH1 β subunit in the other (Fig.1 A-C). This interaction would explain why mutation of either Arg49 or Arg53 to Lys disrupts dimer formation [47] as this would prevent the interaction between SQDG and PufX that appears to hold the two monomers together.

**Figure 1.**
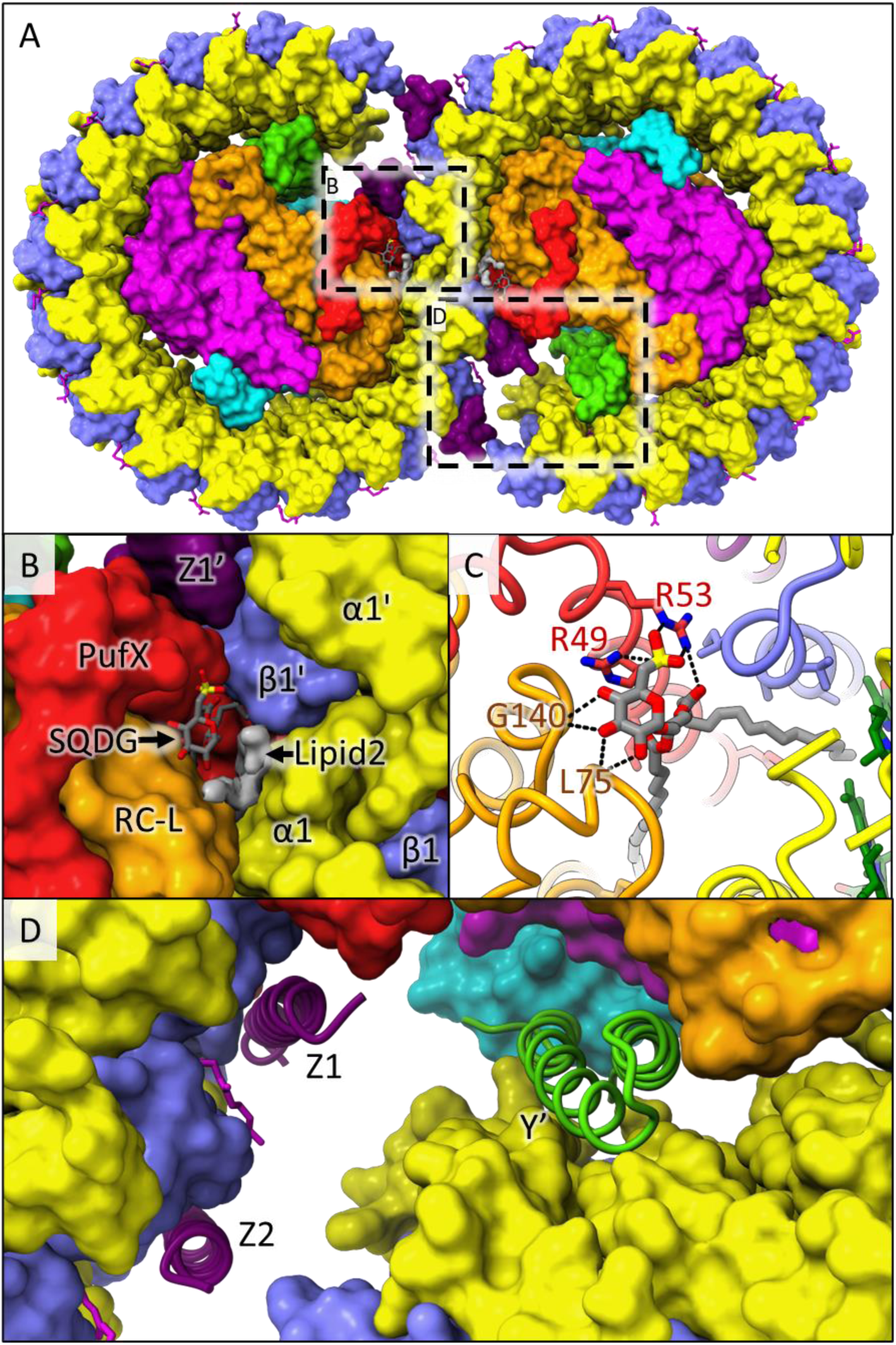
Structure of the dimeric *Rba. sphaeroides* RC-LH1 core complex. (A) Surface view of the complete complex from the lumenal (periplasmic) face. The RC subunits are shown in orange (RC-L), magenta (RC-M) and cyan (RC-H). LH1 subunits are in yellow (α) and blue (β). Additional subunits are in red (PufX), green (protein-Y) and purple (protein-Z). Lipids and cofactors are shown in stick representation in green (BChl), magenta (carotenoids) and grey (SQDG). Unassigned density for lipid 2 is shown as a grey surface. Boxes illustrate areas enlarged in panels B, C and D. (B) Enlarged view highlighting one of the SQDGs and lipid 2 bound at the dimer interface. The subunits from the left monomer (PufX, RC-L, α1 and β1), and those from the right monomer (α1’, β1’ and Z1’) are labelled. (C) A further enlarged view of the SQDG lipid with the protein in ribbon representation. Hydrogen bonds between the SQDG head group and PufX Arg49 and Arg53, and the backbone of RC-L residues Leu75 and Gly140 are shown. (D) Enlarged view of the two protein-Z subunits bound to the left monomer (Z1 and Z2, purple) and protein-Y of the right monomer (Y’, green) in ribbon representation. The rest of the protein is in surface representation.

In this study we investigate the roles of SQDG, ornithine lipids, and the newly identified Y and Z subunits of the RC-LH1 complex by generating a series of mutants deficient in synthesis of specific lipids or lacking the genes that encode protein-Y or protein-Z. The effects of these mutations on RC-LH1 dimerization, RC activity and photoheterotrophic growth have been characterised, providing new insights into these recently identified components of the RC-LH1 core complex.

## Materials and Methods

### Generation of strains and plasmids

The strains used in this study are detailed in Table 1, primer sequences are provided in Table S1, and plasmid information is given in Table S2. Genomic modifications were made using the pK18mob*sacB* plasmid as previously described [48]. Briefly, the target genes were deleted by amplifying ∼400 bp regions upstream and downstream of each gene and joining them by overlap extension PCR, yielding a sequence lacking most of the coding region but leaving the start and stop codons intact and in frame to guard against interference with downstream genes upon genomic modification. The resulting PCR fragments were digested with EcoRI and HindIII and ligated into pK18mob*sacB* cut with the same restriction enzymes. The resulting sequence-verified plasmids were transformed into *E. coli* S17-1 and then conjugated into either wild-type or Δ*crtA Rba. sphaeroides.* Correctly modified strains were isolated following sequential selection with kanamycin (30 µg/ml) and counter-selection with sucrose (10% w/v) and the modified loci were verified by PCR and automated Sanger sequencing (Eurofins Genomics).

**Table 1.** Bacterial strains used in this study.

| Species | Description | Source/reference |
| --- | --- | --- |
| ***Escherichia coli*** |  |  |
| JM109 | Cloning strain for generating plasmid constructs | Promega, UK |
| S17-1 | Conjugative strain for transfer of plasmids to *Rba.*<br>*sphaeroides* | Simon et al 1983 |
| ***Rhodobacter sphaeroides*** |  |  |
| Wild type | Strain 2.4.1 | S. Kaplan* |
| $\Delta crtA$ | Unmarked deletion of *crtA* (Rsp_0272); carotenoid<br>pathway truncated at spheroidene | Chi et al 2015 [49] |
| $\Delta puyA$ | Unmarked deletion of *puyA* (Rsp_7571); does not<br>produce protein-Y | This study; Qian et al<br>2021a [7] |
| $\Delta puzA$ | Unmarked deletion of *puzA* (orf located within<br>Rsp_2385 gene); does not produce protein-Z | This study; Qian et al<br>2021b [8] |
| $\Delta olsBA$ | Unmarked deletion of *olsBA* (Rsp_3826-3827); does<br>not make ornithine lipids | This study; Aygun-Sunar et al 2006 [50] |
| *ΔsqdB* | Unmarked deletion of *sqdB* (Rsp_2569); does not make SQDG | This study; Benning and Somerville 1992 [51] |
| *ΔcrtA ΔpuyA* | Unmarked deletion of *puyA* from *ΔcrtA* background | This study |
| *ΔcrtA ΔpuzA* | Unmarked deletion of *puzA* from *ΔcrtA* background | This study |
| *ΔcrtA ΔolsBA* | Unmarked deletion of *olsBA* from *ΔcrtA* background | This study |
| *ΔcrtA ΔsqdB* | Unmarked deletion of *sqdB* from *ΔcrtA* background | This study |
| *ΔcycA ΔcycI* | Unmarked deletion of *cycA* (Rsp_0296) and *cycI* (Rsp_2577); does not produce cytochrome *c*<sub>2</sub> (CycA) or isocytochrome *c*<sub>2</sub> (Cycl) | This study |

\*Department of Microbiology and Molecular Genetics, The University of Texas Medical School, Houston, TX 77030, U.S.A.

Expression plasmids were generated by amplifying fragments containing *sqdB* or *sqdBDC* with HindIII and BcuI restriction sites. The digested PCR products were ligated into a modified pBBRBB-P*puf*_843-1200_ plasmid in which the *puf* promoter-DsRED fragment was replaced with the *pucBAC* genes and 364 bp upstream of *pucB,* corresponding to the *puc* promoter (P*puc*). A HindIII site was engineered immediately downstream of P*puc* such that *pucBAC* could be replaced with a gene of interest by HindIII-BcuI digestion. The *sqdB* and *sqdBDC* plasmids were conjugated into the Δ*sqdB* strain with selection on kanamycin-containing plates, followed by screening of the genomic *sqdB* locus and the gene(s) in the plasmid by PCR.

### Cell growth and preparation of intracytoplasmic membranes

*Rba. sphaeroides* cells were grown photoheterotrophically in 1 L Roux bottles containing M22 medium [52] under ∼50 µmol m^-2^ s^-1^ illumination from 70 W Phillips Halogen Classic bulbs until they reached stationary phase (an optical density at 680 nm [OD_680_] of ∼3). Strains containing pBBRBB plasmids were supplemented with 30 µg ml^-1^ kanamycin. Cells were harvested by centrifugation at 4,000 RCF for 30 min at 4 °C then resuspended in 5 mL 20 mM Tris-HCl pH 8. Following addition of a few crystals of DNaseI and lysozyme, cells were disrupted via two passes through a chilled French press at 18,000 psi followed by removal of unbroken cells and insoluble debris by centrifugation at 25,000 RCF for 30 min at 4 °C. The supernatant was layered on top of 15/40% (w/v) discontinuous sucrose gradients and centrifuged at 85,000 RCF in a Beckman Type 45 Ti rotor at 4°C for 10 h. A pigmented band of ICM formed at the 15/40% interface, which was harvested using a serological pipette and stored at -20°C until required.

### Thin-layer chromatography (TLC) of lipids

TLC was performed based on a method by Swainsbury et al [17] with some modifications. TLC plates were activated by soaking in 0.15 M ammonium sulphate for 15 min and placed in an oven at 160 °C for1 h. 30 µl volumes of membrane fractions, at an optical density at 875 nm (OD_875_) of 10 were dissolved in 100 µL of 50:50 methanol:chloroform, centrifuged in a benchtop microcentrifuge at 16,000 RCF for 5 min and the lower chloroform phase was removed. 5-10 µl of each lipid standard (∼5 mg mL^-1^ in chloroform) and membrane samples were loaded on to the TLC plate and run in 85:15:10:3.5 chloroform:methanol:acetic acid:water for 45 min. The plate was dried for 5 min before being submerged in 50% (v/v) H_2_SO_4_ for 10-20 s and dried, prior to heating at 160 °C for 10 min to develop the lipid bands.

### Fractionation of photosynthetic complexes by rate-zonal centrifugation

Membranes harvested from discontinuous sucrose gradients were diluted at least 5-fold in 20 mM Tris, pH 8 and pelleted at 185,000 RCF for 2 h using a Beckman Type 45 Ti rotor at 4 °C. Pelleted membranes were resuspended in approximately 100–200 μL of 20 mM Tris pH 8. Six OD_875_ units of resuspended membranes were solubilised in 3% (w/w) n-dodecyl-β-D-maltoside (β-DDM) in a total volume of 375 μl for 1 h at room temperature before centrifugation at 15,000 rpm at 4 °C for 1 h in a microcentrifuge. The supernatant was collected and layered on top of discontinuous sucrose gradients containing steps of 20, 21.25, 22.5, 23.75 and 25% (w/w) sucrose in 20 mM Tris-HCL pH8 and 0.03% (w/v) β-DDM. Gradients were centrifuged in a Beckman SW41 Ti rotor at 125,000 RCF for 40 h at 4 °C. Each gradient was performed in technical triplicate from two biological repeats. Pigmented bands were harvested with a peristaltic pump for downstream processing. If being used for turnover assays, RC-LH1 monomer and dimer bands were buffer exchanged into 50 mM Tris at pH 7.5 with 100 mM NaCl and 0.03% w/v β-DDM by spin concentration with 50,000 MWCO centrifugal concentrators (Sartorius).

To attempt to reform RC-LH1 dimers from monomers of the Δ*sqdB* strain by provision of SQDG, the monomer band harvested from a discontinuous sucrose gradient of the Δ*sqdB* strain was concentrated to 500 µL in a 50,000 MWCO centrifugal concentrator (Sartorius). This sample was split and incubated with and without 0.05% (w/v) SQDG for 24 h before application to another discontinuous sucrose gradient, as described above.

### Bioinformatics

Homologues of PufX, protein-Y, protein-Z and the SQDG producing enzyme SqdB were identified by performing tblastn (protein to translated nucleotide) searches against the whole genome contigs (wgs) database on the NCBI BLAST webserver (https://blast.ncbi.nlm.nih.gov/Blast.cgi). Search parameters were calibrated using PufX as a benchmark, which is known to be highly divergent, and were complicated by its short sequence. The settings used to provide positive hits to all PufX proteins in the target species were: Expect threshold = 20; Word size = 2; Max matches in a query range = 0 (default); Matrix BLOSUM62 (default); Gap Costs = Existence: 11, Extension: 1 (default); Compositional adjustments = No Adjustment; Filter Low complexity regions = off. Searches were performed using the *Rba. sphaeroides* PufX sequence as a template (UniProt entry P13402) against each species individually. Top scoring hits for each species were verified by viewing relevant UniProt entries for each sequence, which were annotated as PufX in all cases. Next, searches for LH2 α (UniProt entry Q3J144), protein-Y (UniProt entry U5NME9), the resolved sequence of protein-Z extracted from PDB entry 7PQD, and SqdB (UniProt entry Q3J3A8) were performed using the same settings as for PufX. The top scoring hits from each species were verified by manually inspecting the genomes in the KEGG database (https://www.genome.jp/kegg/) or GENBANK, rejecting sequences embedded within larger genes or in unrelated genomic regions. LH2 hits were also verified via previous experimental determination of LH2 production in these species. A phylogenetic tree of 16S rRNA sequences was generated via a nucleotide BLAST of the *Rba. sphaeroides* 16S RNA (Genbank entry KF791043.1) against the whole genome contigs (wgs) database for all species analysed in this study, and the non-photosynthetic alphaproteobacterium *Caulobacter vibrioides* as an outgroup to root the tree. The tree was rendered using FigTree v1.44 (available at http://tree.bio.ed.ac.uk/software/figtree/) following flipping of some nodes to sort species by whether they form RC-LH1 dimers. We noted that *puyA* was located near to *otsB* (encoding trehalose-phosphatase) in the genome of the species that have this gene. Whole genome shotgun sequences of *Rba. changlensis* were searched manually for an ORF resembling *puyA* in the vicinity of *otsB.* The sequence in supplementary Figure 6 was found in *Rba. changlensis* strain DSM 18774 NCBI Reference Sequence: NZ_QKZS01000001.1.

### Purification of *Rba. sphaeroides* cytochrome *c*_2_

A cytochrome *c*_2_ overproduction strain was constructed by expressing *cycA* under the control of the constitutive P*puf_842-1200_* promoter on the pBBRBB plasmid in a strain lacking the genomic copies of *cycA* and *cycI*. Cell pellets from semi-aerobic cultures (1.6 L of media in 2.5 L conical flasks shaken at 180 rpm at 34 °C) were resuspended in periplasmic extraction buffer (100 mM Tris-HCl pH 8, 500 mM sucrose and 50 mM NaCl) supplemented with a tablet of EDTA-free protease inhibitor (Merck) up to a total volume of 40 mL. 0.8 g of solid sodium deoxycholate (Sigma) was added to the cell resuspension and, after an hour of incubation at 4 °C in the dark, spheroplasts were pelleted at 30,000 RCF for 30 minutes. The supernatant was transferred to a fresh centrifuge tube and 6.25 mL of deoxycholate precipitation solution (1 M Ammonium acetate at pH 5, 250 mM MgSO_4_) added, before the precipitate was removed by an identical centrifugation step. The supernatant from this second spin step was subsequently passed through 2 0.22µm filters (Sartorius) and made up to 500 ml with 50 mM ammonium acetate buffer at pH 5 before loading onto a 30 mL SP Sepharose column (Cytiva). Cation exchange was performed to purify the cytochrome using a gradient of 13 – 23% buffer B (50 mM ammonium acetate, 1M NaCl).

### RC-LH1 turnover assays

Turnover assays were conducted under steady state conditions in a similar fashion to that described in Swainsbury et al., (2021) [23], using 300 µl solutions containing 5 µM reduced cytochrome *c*_2_, 50 µM ubiquinone-2 (Merck) and 0.01-0.05 µM RC-LH1 in a buffer mixture containing 50 mM Tris, 100 mM NaCl, 1 mM sodium-D ascorbate and 0.03 % w/v β-DDM. Following overnight dark adaption, 300 µL of each reaction mixture were placed in a quartz cuvette and monitored at 550 nm using a Cary60 spectrophotometer (Agilent technologies). After 10 seconds, excitation energy was delivered via a fibre optic cable from an 880 nm M810F2 LED (light-emitting diode) (Thorlabs Ltd., UK) driven at 100% intensity using a DC2200 controller (Thorlabs Ltd., UK). The data were processed by fitting the linear initial rate over 0.025-0.1 seconds, starting from the first data point where the absorbance started dropping continuously. Rates were normalised to e^-^/RC/sec by dividing the cyt *c*_2_ oxidation rate per second by the RC-LH1 concentration. The concentrations of the RC-LH1 complexes were determined using an extinction coefficient of 3,000 mM^-1^ [53], except for the monomeric Δ*puyA* complex, which we predicted to have an extinction coefficient of 2835 mM^-1^ based upon spectra normalised to the RC bacteriopheophytin band in Fig.3.

## Results

### Generation and verification of strains lacking SQDG and ornithine lipids

As shown in Fig. 2A, SQDG synthesis from UDP-glucose requires four enzymes: SqdB, SqdA, SqdC and SqdD [51]; we deleted the *sqdB* (Rsp_2569) gene from the *sqdBDC* operon to prevent the first step of SQDG biosynthesis (Fig. 2C). By contrast, just two enzymes, OlsA and OlsB, are required to produce ornithine lipids from L-ornithine [50] (Fig. 2B). The *olsB* (Rsp_3826) and *olsA* (Rsp_3827) genes are arranged into an operon in which the 3′ end of *olsB* and the 5′ end of *olsA* overlap (Fig. 2D); this gene pair was deleted to prevent ornithine lipid production.

**Figure 2.**
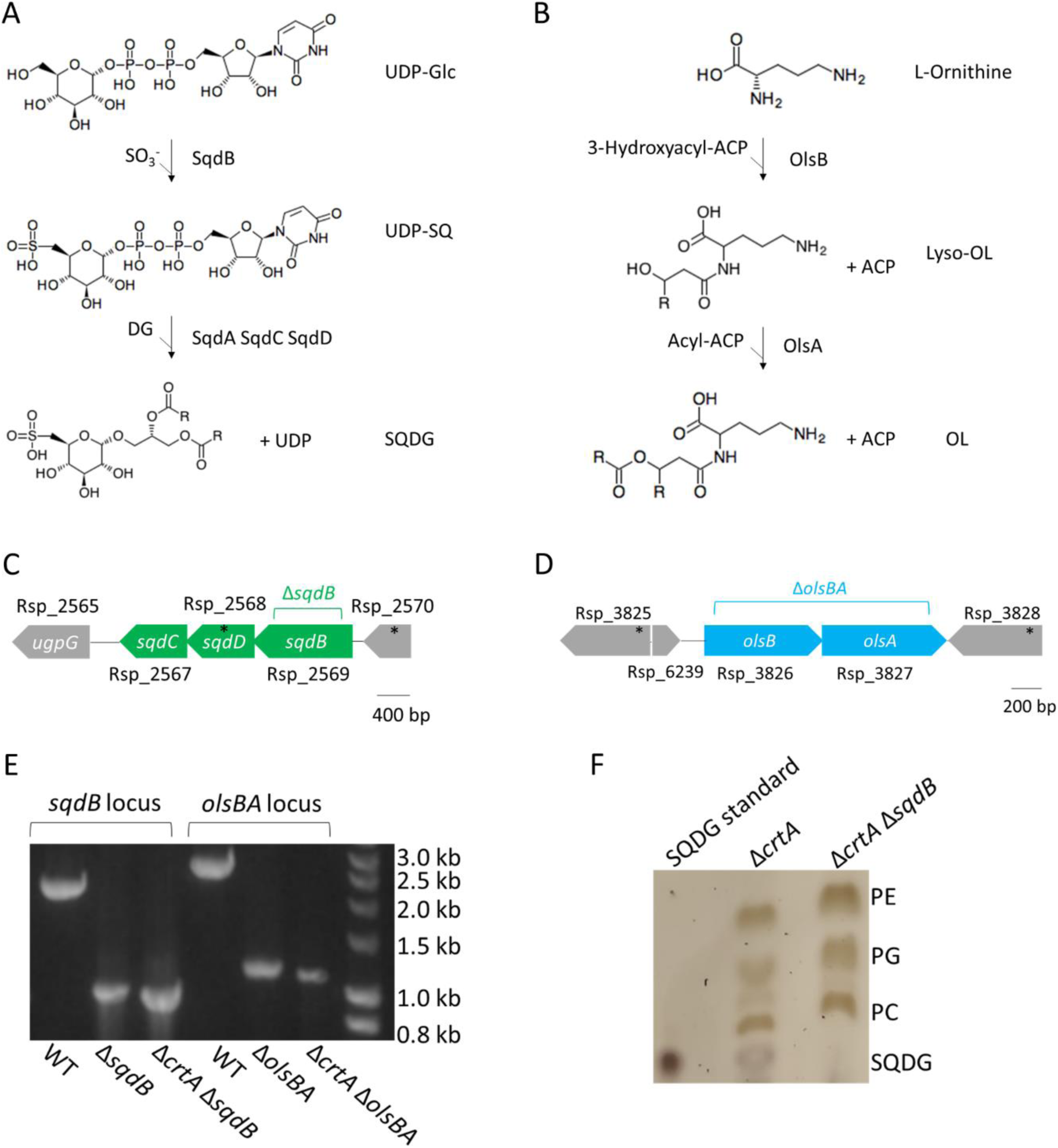
Lipid biosynthesis pathways and deletion of *sqdB* and *olsBA* and PCR verification of knockout strains. (A) sulfoquinovosyl diacylglycerol (SQDG) is synthesised in two steps. The first step is the addition of sulphite to UDP-glucose (UDP-Glc) to produce UDP-sulfoquinovose (UDP-SQ) by SqdB. Next, diacylglycerol is added and UDP is removed by SqdA, C and D to produce SQDG [51]. (B) Ornithine lipids are synthesised by the acyltransferases OlsB and OlsA, which sequentially add a 3-hydroxyacyl group then an acyl group to L-ornithine using 3-hydroxyl-ACP and acyl-ACP as substrates, respectively [54]. (C) Structure of the *sqdBDC* operon. The region labelled Δ*sqdB* (encompassing Rsp_2569) was removed to abolish SQDG biosynthesis. (D) Structure of the *olsBA* operon. The labelled region spanning *olsB* (Rsp_3826) and *olsA* (Rsp_3827) was removed to prevent OL biosynthesis. (E) Agarose gel of ethidium bromide-stained PCR products showing size differences for the amplified regions spanning the *sqdB* and *olsBA* genes showing a clear reduction in size in the knockout strains relative to the wild-type. (F) TLC plate showing loss of SQDG lipid in the Δ*sqdB* strain. A standard for SQDG was run in lane A and a band of the expected size can be seen in samples from Δ*crtA* but not in Δ*crtA* Δ*sqdB* confirming the loss of SQDG biosynthesis.

**Figure 3.**
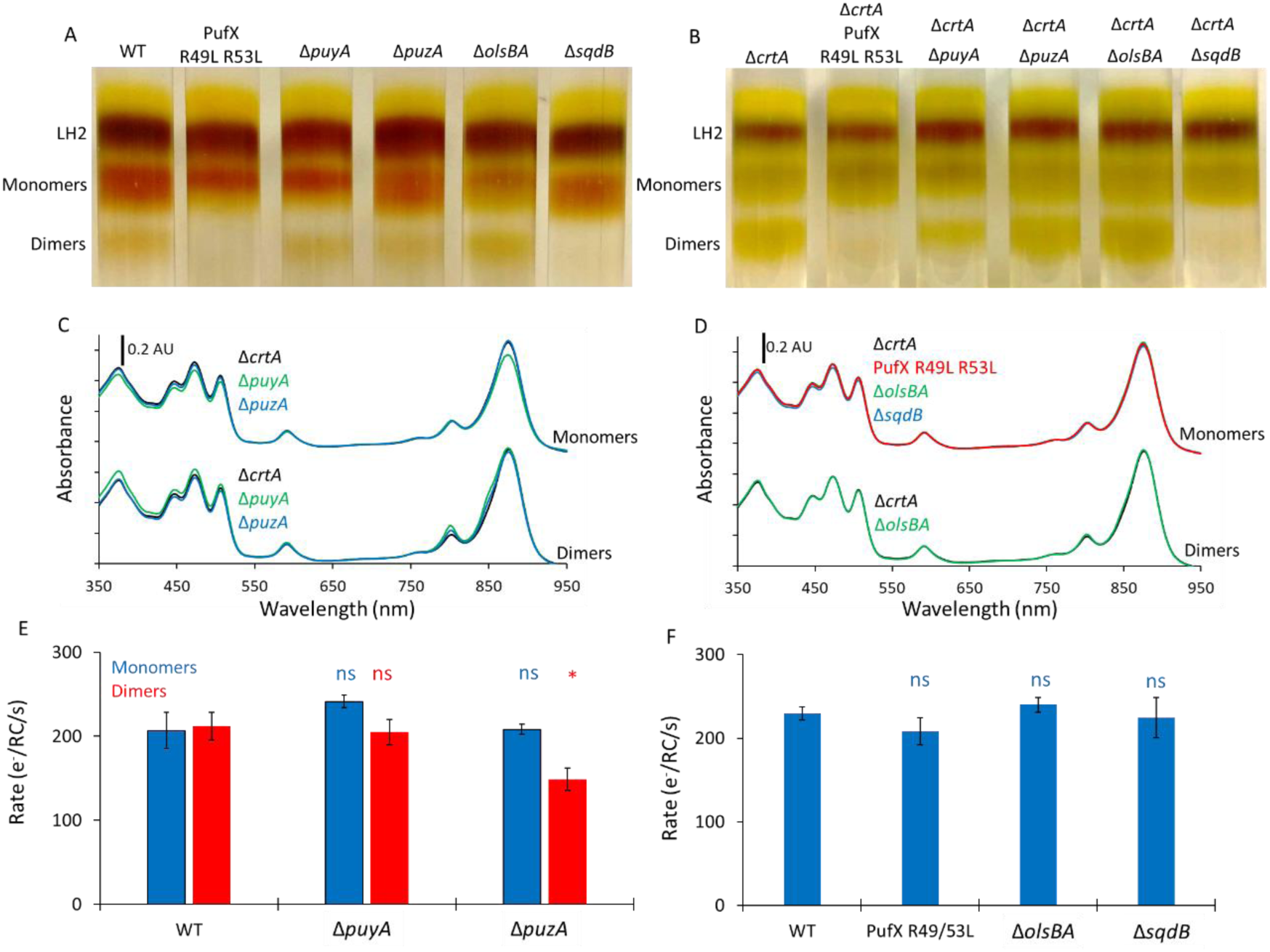
The effects of removing protein-Y, protein-Z, SQDG or ornithine lipids on RC-LH1 dimer formation, absorbance spectra and RC activity. (A-B) Solubilised chromatophore membranes separate into bands of LH2, RC-LH1 monomer and RC-LH1 dimer when centrifuged on sucrose step gradients. Panel A shows the wild-type strain (WT), a control strain that does not produce dimeric RC-LH1 (PufX R49L R53L), a strain lacking protein-Y (Δ*puyA*), a strain lacking protein-Z (Δ*puzA*), a strain that cannot produce ornithine lipids (Δ*olsBA*), and a strain that cannot produce SQDG lipids (Δ*sqdB*). Panel B shows sucrose gradients for the strains in (A) in a background that is also deficient in the *crtA* gene encoding spheroidene monooxygenase (Δ*crtA*). (C-D) UV/Vis/NIR absorbance spectra of the monomer and dimer RC-LH1 bands harvested from the gradients in (B). Panel C shows spectra for the Δ*crtA* strain, and those also harbouring the Δ*puyA* and Δ*puzA* mutations. Panel D shows spectra for the Δ*crtA* strain, and strains also harbouring the RC-LH1 dimer-deficient PufX R49L R53L mutations, and the Δ*olsBA* and Δ*sqdB* genes. (E) Turnover assays for monomeric and dimeric complexes from the WT, and strains lacking proteins Y and Z in strains also lacking CrtA. (F) Turnover assays for the monomeric WT, PufX R49L R53L monomeric control, and monomeric lipid-deficient mutants produced in the wild-type background strain. Rates in E-F show moles of cyt *c*_2_ oxidised per second per mole of RC during illumination using an 810 nm LED of a solution containing 0.01 (E) or 0.05 (F) μM RC-LH1 and 5 μM cyt *c*_2_. T-tests were performed relative to rates for monomeric or dimeric WT complexes where * denotes a p value from 0.01-0.05 and ns denotes a p value > 0.05.

Previous work has shown that the deletion of *crtA* influences the levels of RC-LH1 dimer formation [49]. We therefore deleted *sqdB* and *olsBA* in both a wild-type background and in a strain lacking the spheroidene monooxygenase (Δ*crtA*) to better visualise any effect of lipid content on the monomer-dimer ratio (Fig. 3 A-B). TLC confirmed the loss of SQDG when *sqdB* is deleted (Fig. 2F) but ornithine lipids were not detectable, preventing their analysis using this method. Under photoheterotrophic conditions, growth of the strains unable to produce SQDG or ornithine lipids was indistinguishable from strains with unaltered lipid biosynthesis (see Materials and Methods) (Supplementary Fig. 1), and the UV/Vis/NIR spectra of isolated chromatophore membranes from these strains were also very similar (Supplementary Fig. 2). Therefore, removal of SQDG or ornithine lipids did not result in obvious phenotypes with respect to either growth or spectral features.

### The loss of SQDG prevents the formation of dimeric RC-LH1 core complexes

The removal of SQDG in wild-type and Δ*crtA* strains results in a complete loss of observable dimer formation, whereas the removal of ornithine lipids has no discernible effect upon the monomer to dimer ratio of RC-LH1 complexes (Fig. 3A-B). The absence of SQDG-free RC-LH1 dimers supports the assignment of SQDG in the cryo-EM structure and highlights the essential role of this lipid for RC-LH1 dimer formation in *Rba. sphaeroides*. The absence of dimers from a control strain known to only produce monomers, in which the PufX Arg49 and Arg53 residues that hydrogen-bond the SQDG headgroup are replaced with leucines [7,8,47], further supports the essential role of SQDG in dimer formation. The normal growth of the cells, and near-identical absorbance spectra of complexes from strains unable to produce SQDG and ornithine lipids (Supplementary Fig. 1 and Fig.3 C-D), suggests that RC-LH1 monomers are properly assembled without these lipids. Introducing *sqdB* to the Δ*sqdB* strain on a replicative plasmid restores some formation of dimers, whilst introducing the complete *sqdBDC* operon fully restores dimer formation to WT levels (Supplementary Fig. 5A). To test whether SQDG could induce dimerization of fully assembled monomeric complexes, we attempted to dimerise monomeric RC-LH1 complexes from the Δ*sqdB* strain by the addition of SQDG. The monomer band harvested from a 20-25% discontinuous gradient of Δ*sqdB* membranes was incubated with or without 0.05% w/v SQDG for 24 hrs, but no significant dimer formation was seen (Supplementary Fig.5B). The inability of exogenously added SQDG to induce dimerization suggests that this lipid has to be fully integrated during the *in vivo* assembly pathway, in order to create the wide range of interactions that stabilise the RC-LH1 dimer. These include a series of hydrogen bonds with the backbone of the RC-L subunit, the salt bridge complex with PufX Arg49 and Arg53, and hydrophobic contacts with the transmembrane region of the opposing LH1β1 subunit and the BChl1 macrocycle on the other side of the complex [8].

To test whether SQDG and ornithine lipids affect RC-LH1 activity *in vitro*, we monitored the light-driven oxidation rate of cytochrome *c*_2_ by monomeric and dimeric complexes in the presence of ubiquinone-2 (an analogue of the native substrate, ubiquinone-10, with a shortened isoprene tail). In monomeric RC-LH1 complexes from the wild-type background, there was no significant difference in turnover rates between the Δ*sqdb* and Δ*olsBA* mutants and equivalent complexes with both lipids, demonstrating no impairment of assembly or function in the absence of either lipid (Fig. 3F). Therefore, we can conclude that whilst deletion of *sqdB* and therefore removal of SQDG biosynthesis precludes dimer formation, there is no functional effect on the monomeric complex.

### The absence of proteins Y and Z does not prevent dimer formation

As the *puyA* ORF (Rsp_7571) encoding protein-Y is isolated in the genome, we were able to excise it without disturbing upstream or downstream genes, leaving behind a sequence encoding 6 residues in the genome of the unmarked Δ*puyA* strain. The gene encoding protein-Z is located within the Rsp_2385 open reading frame on chromosome 1 (1014511-1014819) but is transcribed in the opposite direction [8]. Most of this gene was deleted to make a Δ*puzA* strain, with just the sequence encoding 7 residues left intact in the genome.

Rate-zonal centrifugation of solubilised chromatophores from these strains show the removal of neither protein-Y nor protein-Z prevents the formation of dimers (Fig.3. A-B). Absorption spectra of each monomer band shows a small but distinct decrease in absorbance at 873 nm in strains lacking protein-Y (Fig 3. C). This observation agrees with the finding that the absence of protein-Y results in monomers missing some α and β LH1 polypeptides, thus they have fewer BChls per RC [9,10]. The dimer bands all have near-identical UV/Vis/NIR spectra, suggesting all dimers contain the same number of α and β polypeptides. The monomer to dimer ratio is similar to wild type in strains lacking either protein-Y or protein-Z, which appear to have no significant role in dimer formation.

Activity assays show that the rate of cytochrome *c*_2_ oxidation by wild-type RC-LH1 monomers and dimers are almost identical, and that rates for both oligomeric forms of the Δ*puyA* RC-LH1 complex are similar, even when accounting for the slight reduction of absorbance at 873 nm (Fig 3E). The activity of the monomeric complexes from the Δ*puzA* strain were found to be similar to the monomeric wild-type complexes, but the activity of the dimeric complexes lacking protein-Z was slightly reduced (p = 0.03), suggesting it may have a small, but not essential, functional role exclusive to the dimer (Fig. 3E).

### SQDG, PufX, protein-Y and protein-Z in other *Rhodobacter* species

Aligning PufX sequences shows that the two arginine residues that bind SQDG, Arg49 and Arg53, are universally conserved amongst the *Rhodobacter* species within the *Cereibacter* subgroup (Fig. 5). All of these species also contain the genes for SQDG biosynthesis (*SqdBDC)*, suggesting RC-LH1 complexes in these species may form dimers with PufX and SQDG in a similar way to *Rba. sphaeroides*. It has been observed that *Rba. azotoformans* and *Rba. changelensis* form dimeric RC-LH1 complexes, but this has yet to be verified for other members of the *Cereibacter* sub-group [55]. The Arg residues are not conserved beyond the *Cereibacter* group and a BLAST search for either *sqdB*, *C* or *D* in the species lacking the PufX Arg residues found no results. We would not expect species with monomeric RC-LH1 complexes to require SQDG, but it is interesting to note that there are two species with dimeric RC-LH1 (*Rhodobaca bogoriensis* [56] and *Rba. blasticus* [57]) that contain neither the Arg49 and Arg53 residues in PufX nor the genes for SQDG biosynthesis. Further exploration is required to see if a different lipid, perhaps with a different mode of binding to PufX, is fulfilling the role of SQDG in dimerization.

We identified sequences for protein-Y in all species within the *Cereibacter* subgroup, and they have extremely high sequence homology that exceeds 94%, except for *Rba. changlensis*, where this value is only 38% (see Fig 4. and Fig. S6 for gene alignments). Protein-Z could also be found in all species in the *Cereibacter* subgroup except for *Rba. changlensis*. The fact that the *Rba. changlensis* PufX sequence is also distinct from the rest of the *Cereibacter* subgroup, and its clear separation in phylogenetic trees generated by its 16S RNA (Fig.4), indicates it may be a more distant relative of *Rba*. *sphaeroides*. We also note that sequence homology was quite high in the first 30 residues of protein-Z, corresponding with the 31 residues resolved in the cryo-EM structure [8], but very low in the rest of the sequence, suggesting function is limited to the N-terminus.

**Figure 4.**
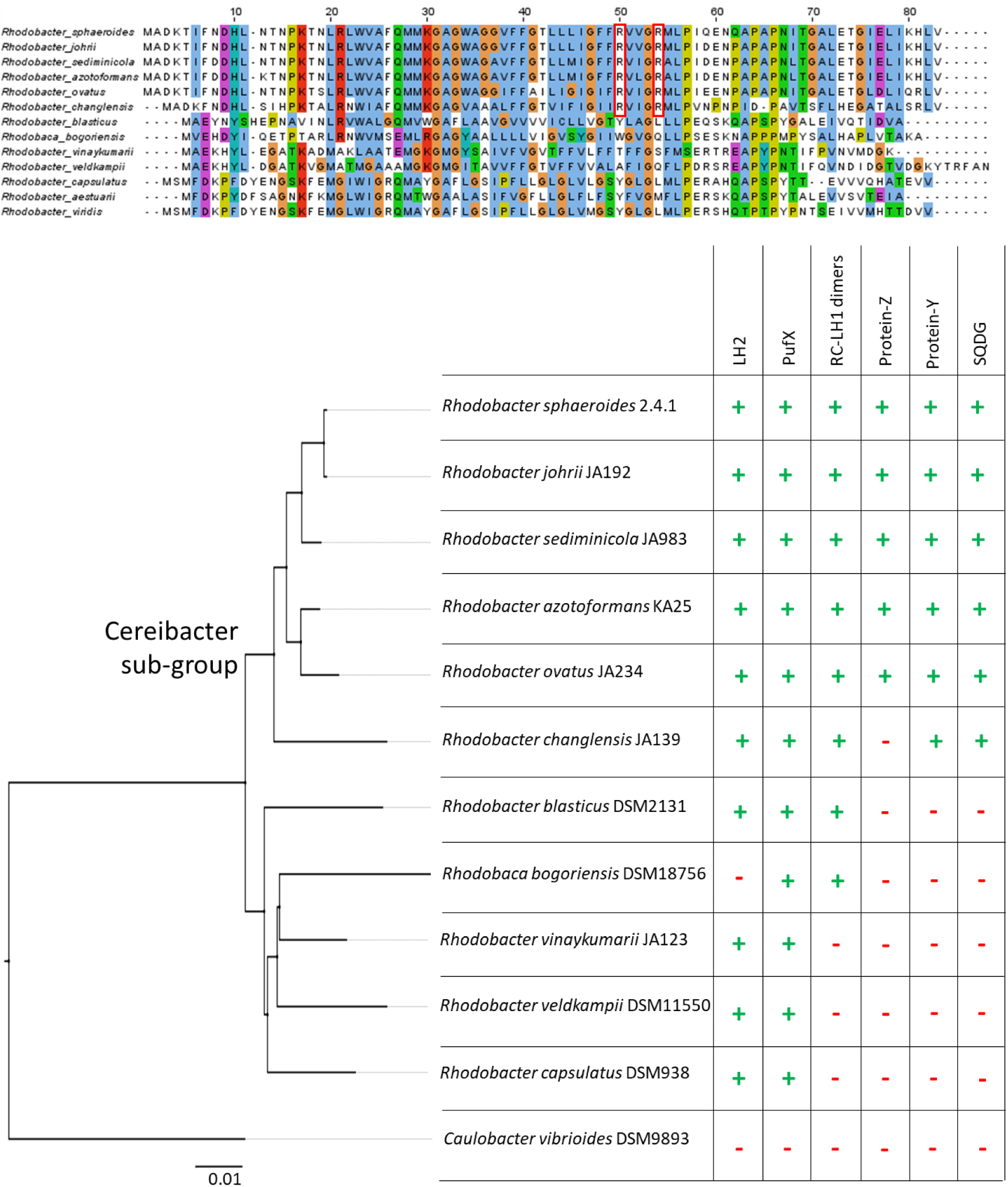
Phylogenetic analysis of the requirements for RC-LH1 dimer formation. (A) Sequence alignment of PufX polypeptides. Red outlined boxes indicate the two arginine residues that bind SQDG in *Rba. sphaeroides*. (B) Left -16S phylogenetic tree of species that produce PufX for *Rhodobacter* species, and other members of the *Rhodobacteriales* order for which the oligomeric state of their RC-LH1 complex is known. The root of the *Cereibacter* subgroup is labelled. *Caulobacter vibrioides*, a non-photosynthetic alphaproteobacterium, is included as an outgroup to root the tree. Right – A table showing the presence or absence of genes encoding the LH2 complex, PufX, protein-Y and protein-Z, whether the species produces dimeric RC-LH1 complexes, and whether it produces SQDG as inferred by the presence of the *sqdB* gene. Green + symbols indicate presence of a feature and red – symbols indicate its absence.

Sequences with homology to *Rba. sphaeroides* protein-Y and protein-Z were not found outside of the *Cereibacter* subgroup. It may be that if proteins are fulfilling the same role, they are not similar enough to be found through searches using sequence homology. This may be the case for the RC-LH1 complex from *Rhodobaca bogoriensis;* structural analysis reveals a protein in a position similar to that adopted by protein-Y in *Rba. sphaeroides* [58], but the DNA sequence identified in the paper bears no homology to *Rba. sphaeroides puyA*.

## Discussion

### The role of PufX, SQDG and OL in RC-LH1 dimer formation and cellular function

PufX has long been known as a unique component of the RC-LH1 complexes of *Rhodobacter* species [24]. In species that produce dimeric forms of the RC-LH1 complex (recently renamed to *Cereibacter* [44]), PufX is essential for mediating the interaction between pairs of RC-LH1 complexes to form the S-shaped dimers [8–10,23,41,59]. As such, PufX has been the subject of intense investigation to understand how and why certain forms of this polypeptide drive RC-LH1 dimer formation whilst others from closely related species do not [55]. Various elements of PufX have been investigated as potential points of contact between monomers, including a conserved GxxxG motif [27,60], the N-terminal 12 amino acids [26], and a pair of Arg residues at positions 49 and 53 (*Rba. sphaeroides* numbering) that are unique to those PufX polypeptides in dimer forming species [47]. It was not until the determination of high-resolution structures of RC-LH1 dimers that the structure of PufX was modelled with sufficient detail to elucidate the interactions that bring the two RC-LH1 monomers together [8–10]. These structures revealed that the N-terminal 15 amino acids were unresolvable, presumably because they are disordered in the mature complex and only required during dimer assembly. Additionally, a short “LWVAF” motif was observed close to the cytoplasmic surface of the membrane that mediates a direct protein-protein interaction between the two PufX proteins. The GxxxG motif plays no role in bringing together opposing PufX polypeptides. The high-resolution structures also revealed that PufX residues 44-67 were found to bind to the lumenal surface of the reaction centre L subunit.

The large distance between Arg49 and Arg53 in one PufX and those in the opposing monomer was a surprising discovery because these residues are known to be essential for dimer formation. In strains where these Arg residues are replaced with other amino acids [47], including the Δ*pufX* mutant strain we used to determine the structure of the monomeric complex [7], the formation of RC-LH1 dimers is abolished. Careful inspection of the cryo-EM map for the RC-LH1 dimer revealed density consistent with an SQDG lipid for which the head group was hydrogen bonded to R49 and R53. One of the hydrocarbon tails extends between PufX and the L-subunit within the RC-LH1 monomer, and the second tail extends across the dimer interface towards the opposing monomer [8] (Fig.1). The specific nature of the headgroup binding, which could not reasonably accommodate an alternative lipid, and the requirement for R49 and R53 for dimer formation led us to speculate that the SQDG lipid itself is essential for dimer formation. In this study, the observation that removing SQDG from *Rba. sphaeroides* membranes via disruption of the biosynthesis pathway confirms our hypothesis.

In contrast to our findings with SQDG, the removal of OL had no observable effect on dimer formation. This suggests that “lipid 2” (Fig.1 and Qian et al [8]) may not be an OL, or that the lipid bound in this position can be readily substituted for an alternative. Because the density was ambiguous, possibly because of more disorder for lipid 2 relative to SQDG, we are still unable to assign it reliably. We also cannot determine whether lipid 2 is required for dimer formation. However, its close association with SQDG and its position in a cavity between RC-LH1 monomers would suggest that the presence of a lipid in this location plays an important role.

Surprisingly, removal of SQDG or OL had no observable effect on photosynthetic growth (Fig. S1) or on the activity of the RC-LH1 complexes (Fig. 3E-F). This is consistent with previous observations that disruption of dimer formation does not produce a discernible phenotype under laboratory conditions [47]. This finding also suggests that specific protein-SQDG or protein-OL interactions are not essential in other membrane protein complexes required for growth under photoheterotrophic conditions, although they may be important under other growth modes, as is the case for OL-deficient strains of *Rba. capsulatus* [50]. SQDG is abundant in whole chromatophore membranes and in the annular lipids of complexes extracted using styrene maleic-acid copolymer nanodiscs [17,61,62], so it appears that bound SQDG or OL can often be substituted with other lipids, with the clear exception of the SQDG at the RC-LH1 dimer interface. This is not the first example of essential, ordered lipids being present in RC-LH1 complexes. In most purple bacteria, RCs are known to bind a conserved cardiolipin, the disruption of which via mutation of interacting residues adversely effects RC thermostability [63–65]. There are also many lipids observed between the RC and LH1 ring, many of which are common to all structurally resolved RC-LH1 complexes and interact via conserved residues on the RC and LH1 α subunits [13, 32]. Therefore, this study and those mentioned above highlight the important role of protein-lipid interactions, which are often unresolved in reported RC-LH1 structures.

### The role of proteins Y and Z in dimer formation and RC turnover

It had long been assumed that the RC-LH1 complexes of *Rba. sphaeroides* contained five unique proteins (RC-L, RC-M, RC-H, PufX, LH1 α and LH1 β), so the discovery of proteins-Y and -Z in the cryo-EM structures was unexpected [7–10]. Protein-Y was annotated as a hypothetical protein in the UniprotKB, whist protein-Z was unannotated. Our immediate question was whether these newly discovered components of the complex are required for RC-LH1 dimer formation and photosynthetic growth of *Rba. sphaeroides.* Removal of protein-Z had no observable effect on dimer formation or on photoheterotrophic growth but did slightly lower the rates of *in vitro* cytochrome *c*_2_ oxidation by the dimeric complex (p=0.03) (Fig.3B, Fig.S1 and Fig.3E). We note that these assays were only performed in triplicate under a single condition and so are not exhaustive, with further studies required to elucidate the full impact of removing protein-Z. However, these are beyond the scope of the current study.

The removal of protein-Y does not inhibit dimer formation, as indicated by the similar relative abundance of the monomeric and dimeric complexes (Fig. 3A,B). However, the spectra of the monomeric complex show a decrease of LH1 absorbance at 875 nm relative to RC absorbance at 803 and 760 nm, suggesting a lowered LH1 antenna size. Cao *et al* and Tani *et al* recently determined the structure of RC-LH1 lacking protein-Y and found that, whist both monomer and dimer complexes still form, the final subunits of the LH1 antenna fail to assemble or are dissociated during protein purification [9,10]. Our findings are consistent with the generation of the same complexes with a smaller LH1 antenna. Because protein-Y is distant from the dimer interface and does not interact with PufX or SQDG, it is reasonable that its removal does not affect the ability of the complex to form dimers. Photoheterotrophic growth of the strains unable to produce protein-Y was similar to the wild type, which suggests that the loss of four LH1 subunits has a negligible effect on light-harvesting under our laboratory conditions (Supplementary Fig 1). We previously suspected that protein-Y serves to maintain a channel for efficient quinone exchange, which seems at-odds with our *in vitro* assays. However, loss of the terminal LH1 subunits creates a larger opening in the LH1 ring, which may compensate for the loss of protein-Y at the expense of light-harvesting capacity. Such a loss of capacity would likely not influence growth rates under the illumination conditions used for our experiments.

In strains that lack protein-Z, activity of monomeric complexes was similar to the wild type, which was expected because monomeric RC-LH1 does not bind protein-Z [8]. However, the dimeric complexes that lack protein-Z had slightly lowered activity when compared to wild type dimers (Fig.3E). This suggests that protein-Z may act to ensure the quinone exit channel at the dimer interface is kept open by preventing the complex reverting to the closed state observed by Cao *et al* [9]. Despite the small reduction in activity in purified RC-LH1 dimers lacking protein-Z we could not find an effect on growth rates under our laboratory growth conditions (Supplementary Fig 1).

### The evolution of RC-LH1 dimers is synergistic with the presence of SQDG, protein-Y and protein-Z

To further elucidate the roles of SQDG, protein-Y and protein-Z in RC-LH1 dimer formation, we investigated the genomes of other bacteria capable of forming dimeric RC-LH1 complexes to see if they do so via the same mechanism. By searching the relevant databases, we found that the species reclassified into the *Cereibacter* group, all of which are either known to form dimers or we predict will form dimers, appear to contain the genes for SQDG biosynthesis, PufX with SQDG binding residues, protein-Y, and most contain protein-Z, whereas the *Rhodobacter* species that produce monomeric complexes do not appear to contain any of these components. Whilst the phylogenetic analysis we performed is not exhaustive, we are able to speculate that the formation of RC-LH1 dimers in the *Cereibacter* subgroup evolved with the ability to synthesise SQDG, and the evolution of a variant of PufX that could bind the lipid head group. Subsequently, protein-Y was recruited to maintain efficient quinone diffusion. Finally, protein-Z was recruited to lock the complete RC-LH1 dimer in its final conformational state. Exceptions to this are the dimeric RC-LH1 complexes from more distantly related *Rhodobaca bogoriensis* and *Rba. blasticus,* which cannot produce SQDG and appear to be more closely related to the monomer-producing *Rba. capsulatus* and *Rba. veldkampii* than the *Cereibacter* subgroup. At the time of writing the dimeric structure of the *Rhodobaca bogoriensis* complex is published as a preprint [58] and the coordinates are not available in the PDB. However, unlike *Rba. sphaeroides* its LH1 is composed of 15 LH1 subunits and it does not produce LH2, so it may have achieved RC-LH1 dimer formation by a unique mechanism that warrants further investigation.

## Author contributions

C.N.H, A.H. and D.J.K.S. conceived and supervised the study. E.C.M., A.G.M.B., T.W.R., A.H. and D.J.K.S. performed the research. E.C.M., A.G.M.B., C.N.H., A.H. and D.J.K.S. wrote the manuscript.

---

**Supplementary Figure 1.**
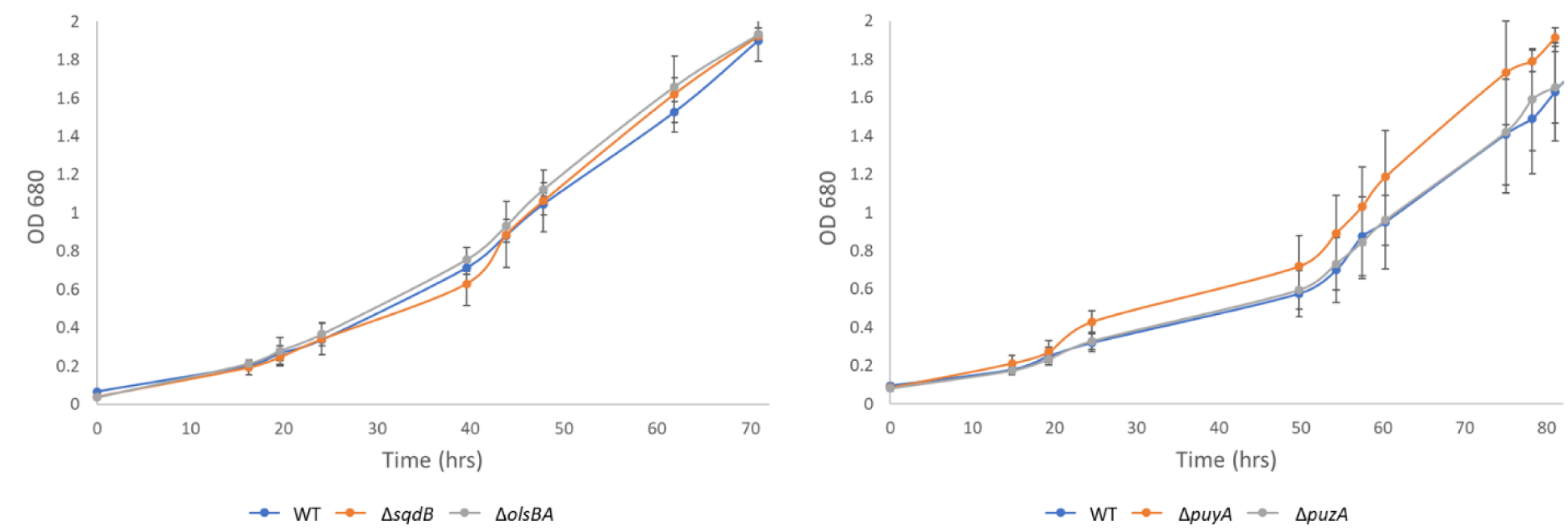
Growth curves of all four knockouts in a WT background used in this study at low light (10umol). No significant phenotype was observed in any strain and all knockouts were confirmed by PCR afterwards. Δ*puyA* showed some variation, but further repeats (data not shown) showed the same growth as WT. A phenotype may yet be apparent at different light intensities.

**Supplementary Figure 2.**
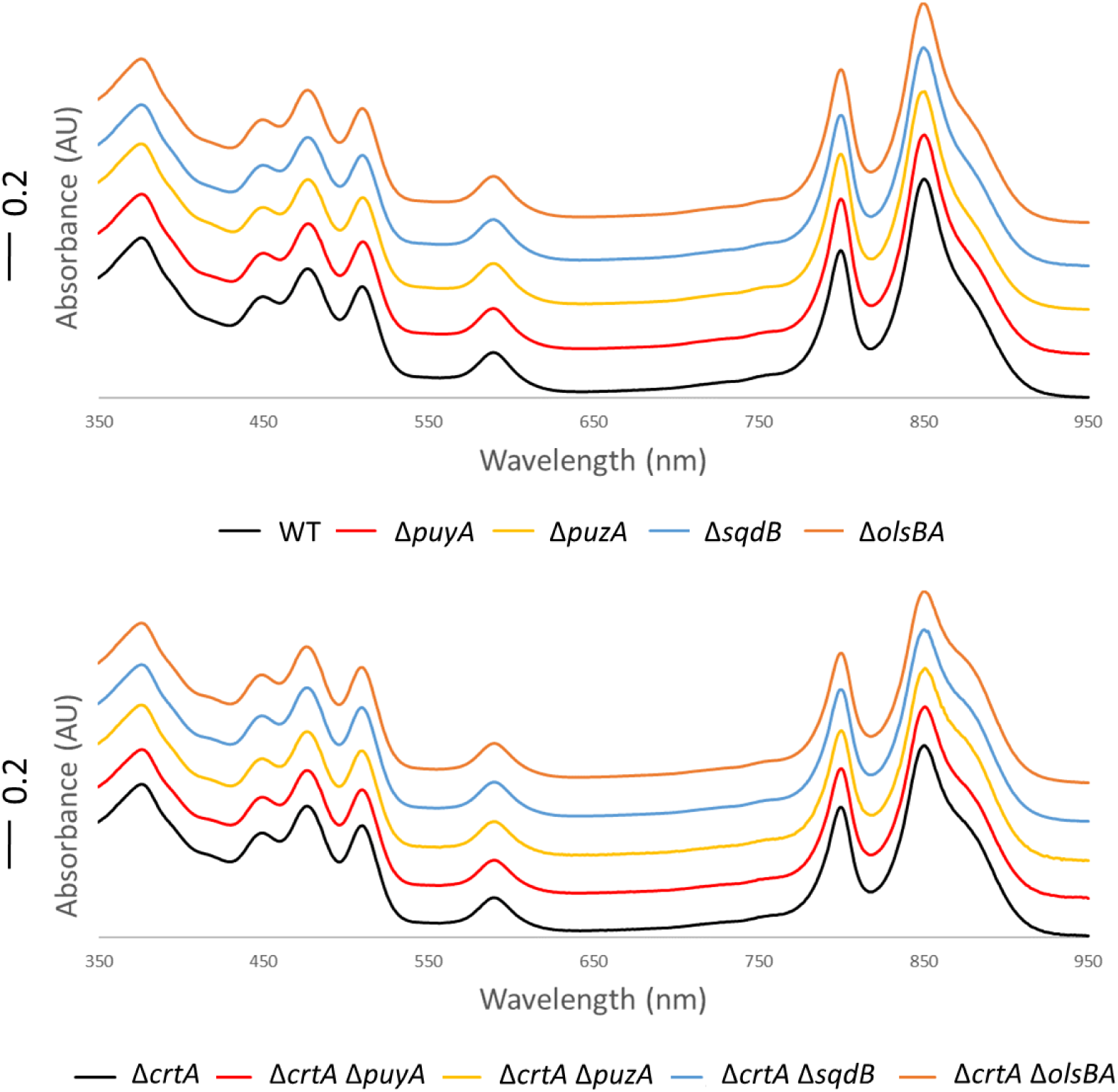
UV/Vis/NIR absorbance spectra of chromatophore membranes from all strains in the WT and Δ*CrtA* backgrounds. Spectra collected of chromatophore membranes isolated from other cellular components by differential centrifugation (see methods). Spectra are offset for clarity.

**Supplementary Figure 3.**
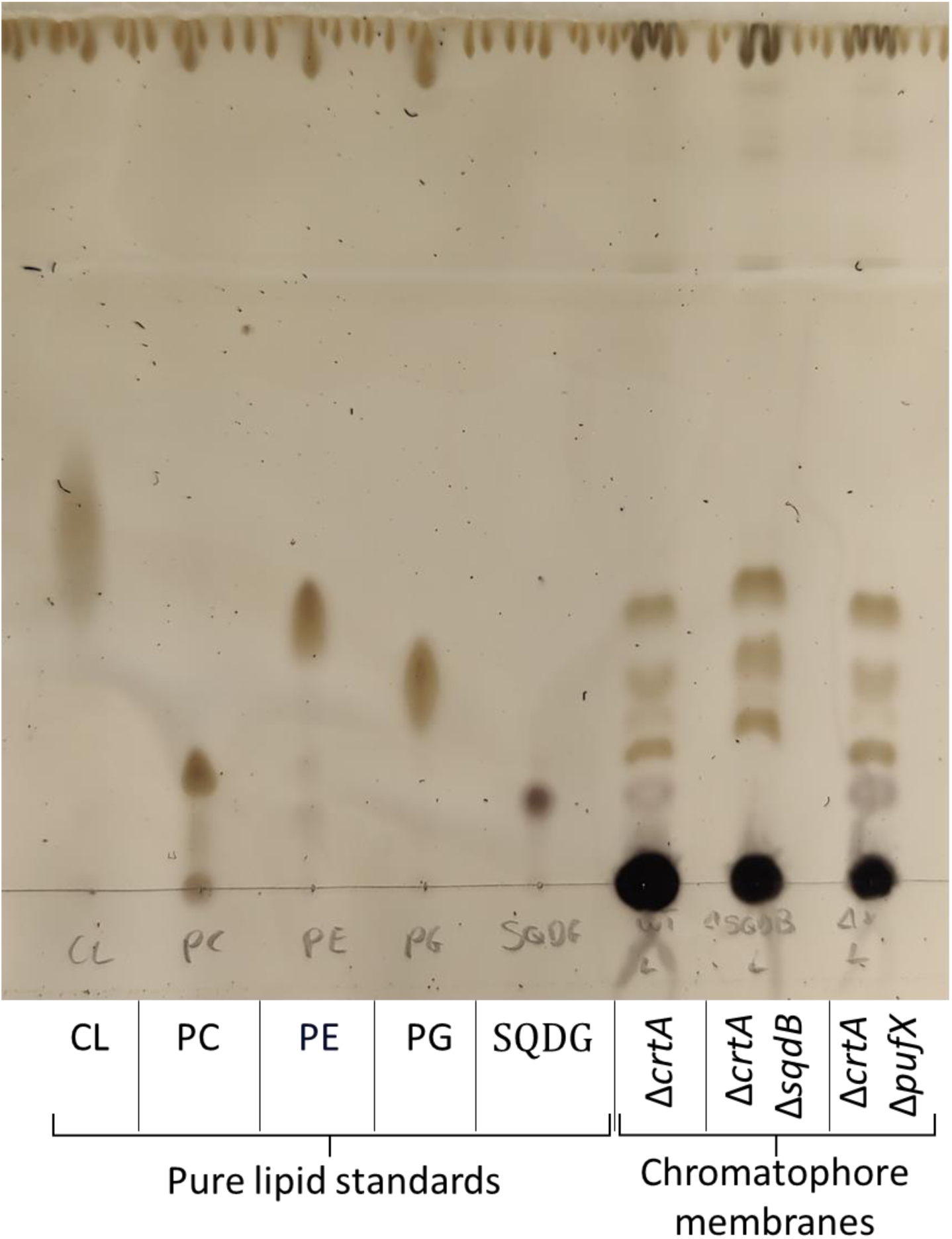
Full TLC plate showing pure lipid standards and lipids extracted from chromatophore membranes. The lipid standards were Cardiolipin (CL), Phosphatidylcholine (PC), Phosphatidylethanolamine (PE), Phosphatidylglycerol (PG), Sulfoquinovosyl diacylglycerol (SQDG). Chromatophore membranes were extracted from *Rba. sphaeroides* cells and isolated by separation on 40/15 % w/v sucrose gradients from the Δ*crtA*, Δ*crtA* Δ*sqdB*, and Δ*crtA* Δ*pufX* strains.

**Supplementary Figure 4.**
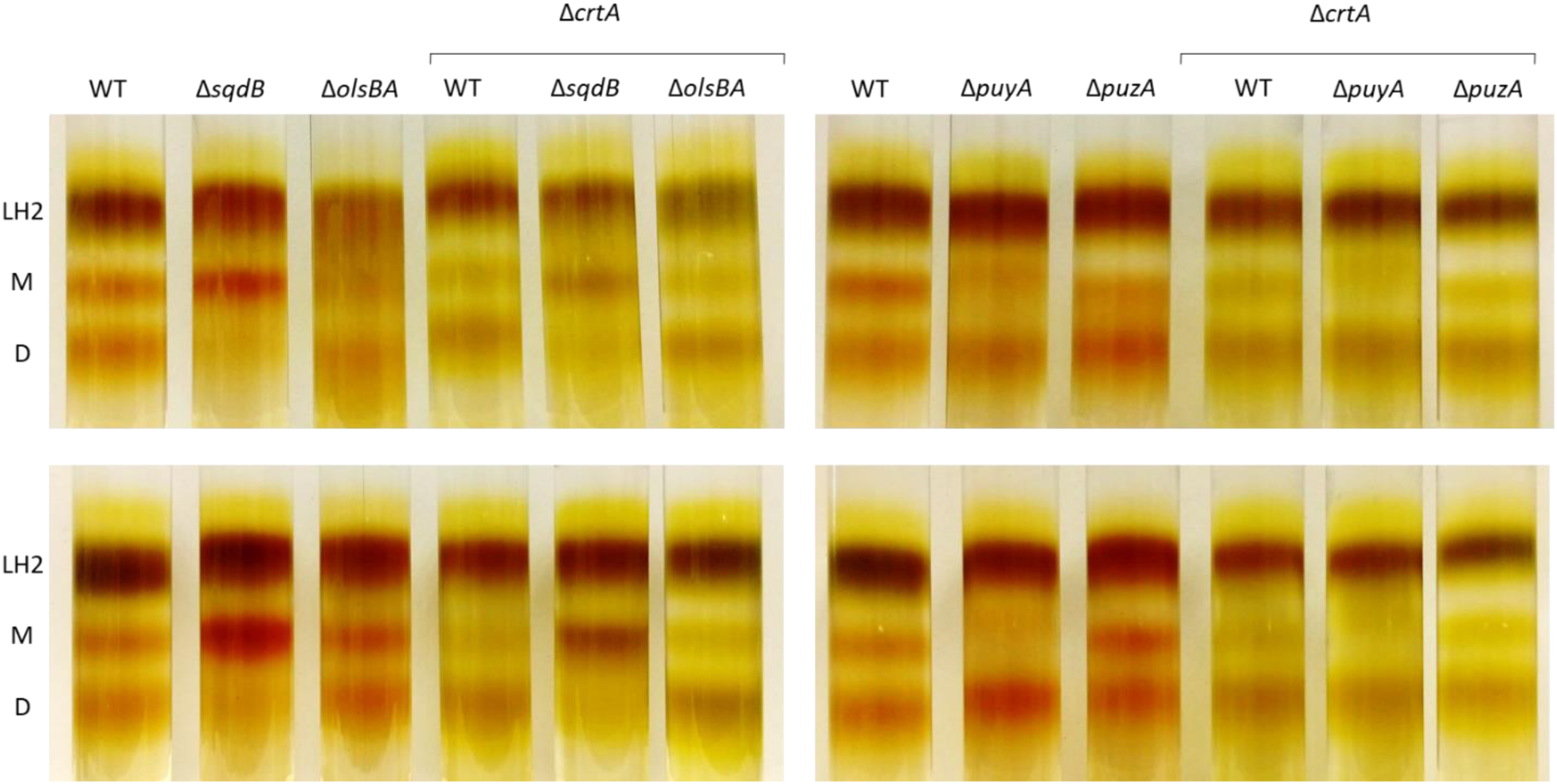
Two further repeats were performed showing the same monomer dimer distribution as presented in Figure 3 of the main paper.

**Supplementary Figure 5.**
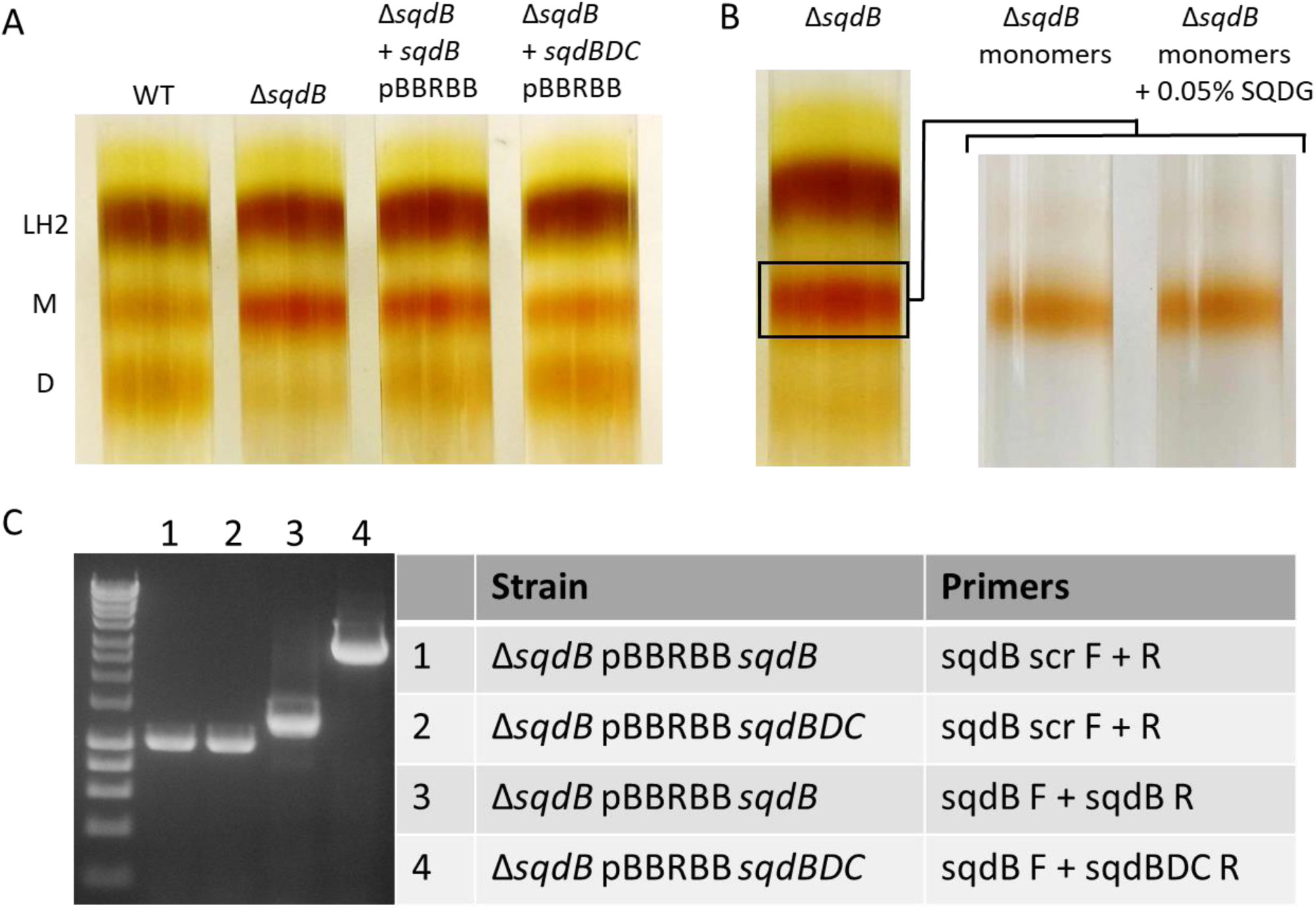
Attempted reconstitution of dimers in the Δ*sqdB* strain by *in trans* complementation or incubation with SQDG. (A) Monomer-dimer gradients of Δ*sqdB* cells expressing *sqdB* from a plasmid show a slight increase in dimer formation. Expression of the *sqdBDC* operon increases dimer expression to WT levels. (B) Purified monomers from Δ*sqdB* incubated with purified SQDG do not spontaneously form dimers. (C) Ethidium bromide-stained PCR products to verify the presence of *sqdB* or *sqdBDC* in pBBRBB-Ppuf_843-1200_ in the Δ*sqdB* background. Lanes 1 and 2 confirm the absence of *sqdB* in the genome of both strains and lanes 3 and 4 confirm the presence of either *sqdB* or *sqdBDC* on pBBRBB.

**Supplementary Figure 6.**
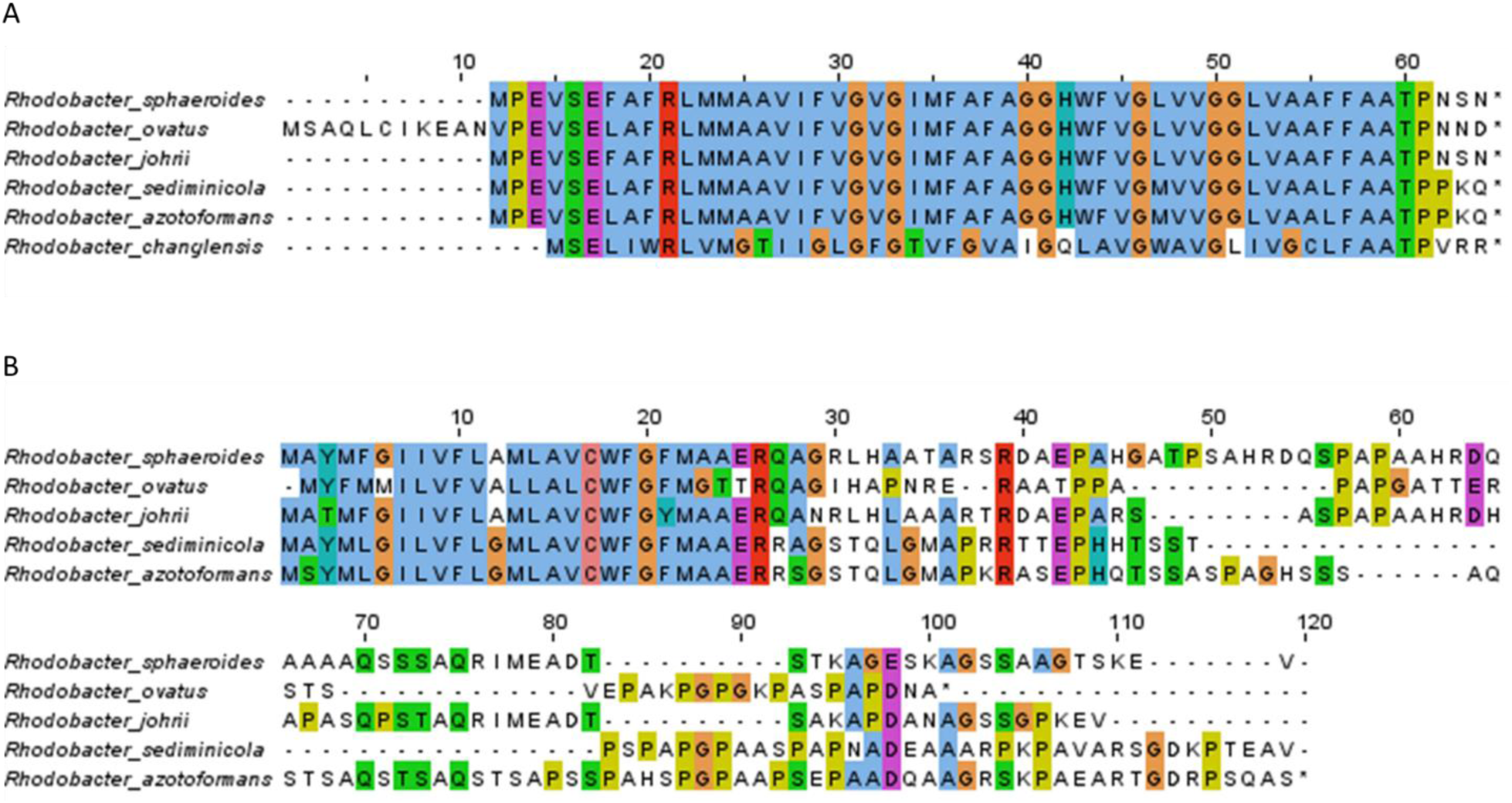
(A) Alignment of sequences for protein-Y from species within the *Cereibacter* subgroup. (B) Alignments for protein-Z. A sequence for *Rba. changlensis* could not be found, potentially due to a lack of homology. With the exception of *Rba. changlensis*, protein-Y shows a very high degree of sequence homology between species. protein-Z has a disordered tail on the C-terminus that is missing in the structure and shows a very high degree of variation between species. Truncations would have to be performed to establish if this region is unnecessary.

**Table S1.**
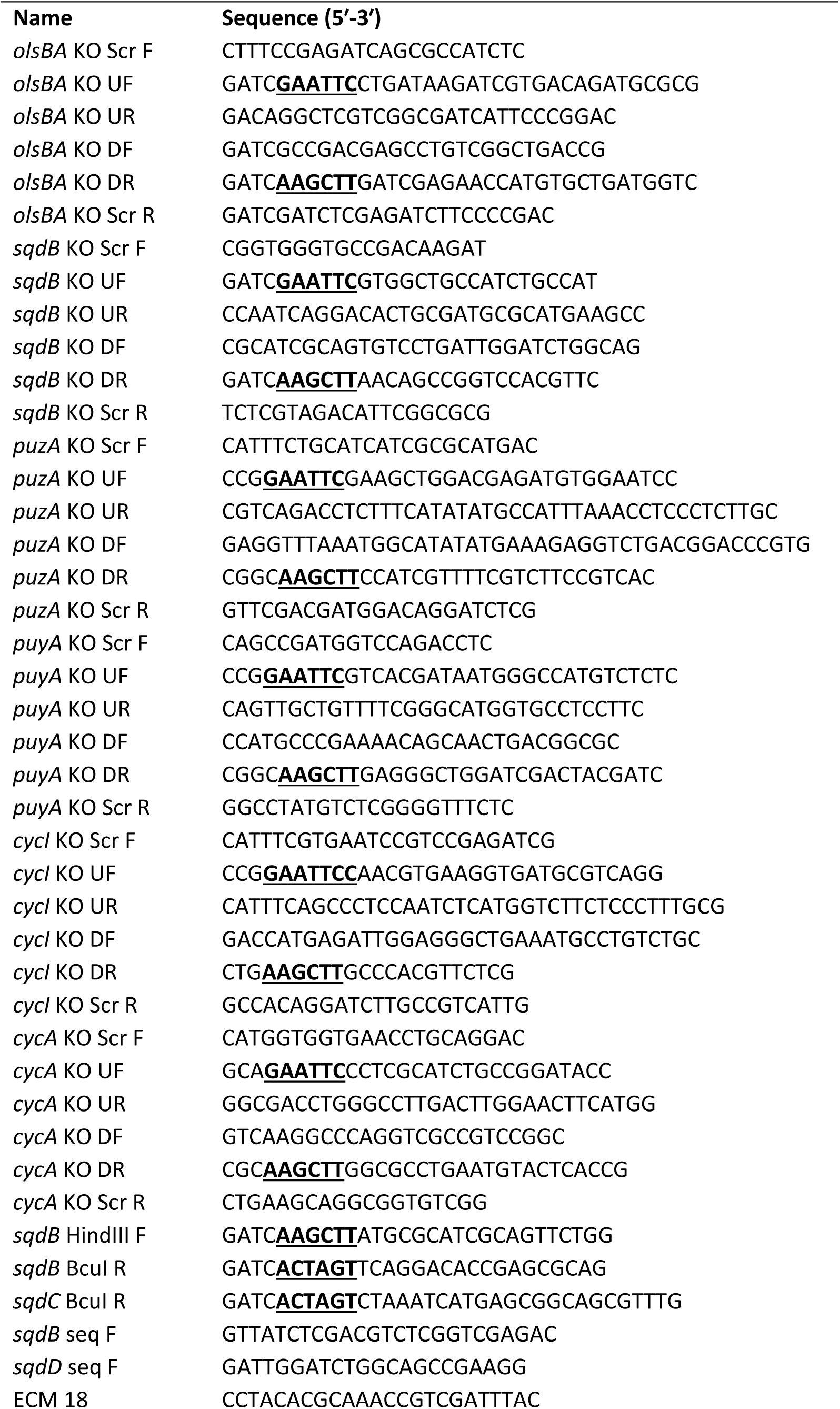
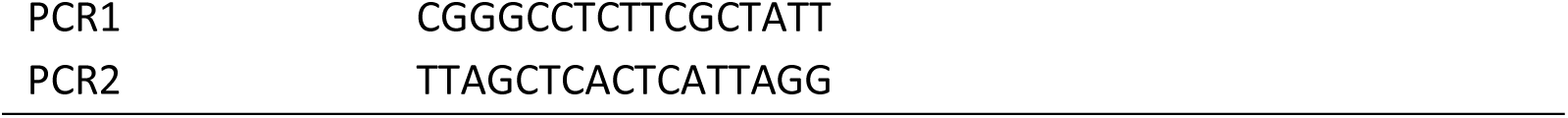
Primers used in this study. Restriction enzyme sites used for cloning are underlined in bold.

**Table S2.** Plasmids used in this study.

| Name | Details | Source/reference |
| --- | --- | --- |
| pk18mobsacB | Allelic exchange vector, Km<sup>R</sup> | Professor J. Armitage* |
| pk18mobsacB::Δ*olsBA* | Construct for unmarked deletion of *olsBA* | This study |
| pk18mobsacB::Δ*sqdB* | Construct for unmarked deletion of *sqdB* | This study |
| pk18mobsacB::Δ*puzA* | Construct for unmarked deletion of *puzA* | This study |
| pk18mobsacB::Δ*puyA* | Construct for unmarked deletion of *puyA* | This study |
| pk18mobsacB::Δ*cycA* | Construct for unmarked deletion of *cycA* | This study |
| pk18mobsacB::Δ*cycl* | Construct for unmarked deletion of *cycl* | This study |
| pBBRBB-Ppuf<sub>843-1200</sub>-DsRed | Replicative expression plasmid, Km<sup>R</sup> | Addgene.org; Tikh et al 2014 |
| pBBRBB-Ppuf<sub>843-1200</sub>-cycA | Plasmid for expression of *cycA* | This study |
| pBBRBB-Ppuc-pucBAC | pBBRBB-Ppuf<sub>843-1200</sub>-DsRed with Ppuf-DsRED replaced with the Ppuc-pucBAC | This study |
| pBBRBB-Ppuc-sqdB | Plasmid for expression of *sqdB* | This study |
| pBBRBB-Ppuc-sqdBDC | Plasmid for expression of *sqdBDC* | This study |

\*Department of Biochemistry, University of Oxford, South Parks Road, Oxford OX1 3QU, U.K.

